# In Silico Defined SARS-CoV2 Epitopes May Not Predict Immunogenicity to COVID19

**DOI:** 10.1101/2021.07.08.451555

**Authors:** Ke Pan, Yulun Chiu, Michelle Chen, Junmei Wang, Ivy Lai, Shailbala Singh, Rebecca Shaw, Cassian Yee

## Abstract

SARS-CoV-2 infections elicit both humoral and cellular immune responses. For the prevention and treatment of COVID19, the disease caused by SARS-CoV-2, it has become increasingly apparent that T cell responses are equally, if not more important than humoral responses in mediating recovery and immune-protection. One of the major challenges in developing T cell-based therapies for infectious and malignant diseases has been the identification of immunogenic epitopes that can elicit a meaningful T cell response. Traditionally, this has been achieved using sophisticated *in silico* methods to predict putative epitopes deduced from binding affinities and consensus data. Our studies find that, in contrast to current dogma, ‘immunodominant’ SARS-CoV-2 peptides defined by such *in silico* methods often fail to elicit T cell responses recognizing naturally presented SARS-CoV-2 epitopes.

## Introduction

Severe acute respiratory syndrome coronavirus 2 (SARS-CoV-2), the highly transmissible respiratory virus responsible for the COVID-19 pandemic outbreak, continues to render significant, lasting impact on global public health and has created an urgent need to develop accurate immunodiagnostics, and effective treatment strategies (1, 2). Rapid dissemination of the SARS-CoV-2 genomic sequence first revealed by Dr. Zhang Yongzhen led to large scale efforts around the world to develop a protective vaccine that could elicit humoral (antibody) and cellular (T cell) responses (3). It follows that the identification of immunogenic epitopes of SARS-CoV-2 recognized by the human immune system would be critical for rational vaccine development.

Using *in silico* prediction algorithms, several investigators have amassed extensive panels of Class I and Class II restricted epitopes to probe SARS-CoV-2-specific T cell responses, in some cases, combining these with overlapping ‘megapools’ spanning regions conserved regions of the genome (4, 5). These peptides have been used to track responses in infected and convalescent individuals (6, 7), design multi-epitope vaccines and used directly or indirectly to measure the breadth and severity of COVID19 disease (7-13). While these studies have uncovered insights into the T cell immunobiology of COVID19, the accuracy of T cell responses using in silico predicted responses and overlapping long peptide (OLP) pools is diminished by a failure to consider whether such epitopes are immunogenic. An immunogenic epitope in this sense is defined as a peptide that is known to be presented by self-MHC, and is capable of eliciting T cells of sufficient affinity that such T cells can recognize target cells endogenously expressing antigen and presenting the antigen-derived peptide in the context of an MHC complex with sufficient surface density as to sensitize the target cell to peptide-specific T cell-mediated recognition. In essence, an immunogenic epitope of SARS-CoV-2 requires both direct sequencing of peptides presented by MHC as well as empiric validation of T cell immunogenicity.

## Results

### SARS-CoV-2 peptides defined ‘*in silico*’, fail to elicit T cells that recognize SARS-CoV-2 antigen expressing targets

As an initial screen of known predicted epitopes for immunogenicity, we selected Class I-restricted peptides to the SARS-CoV-2 Spike protein (SP) and membrane glycoprotein (MGP) based on a literature search of studies where such ‘*in silico*’ predicted peptides were described as ‘immunodominant’. These peptides have previously been reported to be ‘immunodominant’ on the basis of their ability to generate high levels of peptide-specific responses from the PBMC of COVID19+ patients and, surprisingly, in some healthy donors as well (apparently as a result of cross-reactive responses from T cells elicited in the past to non-pathogenic SARS viruses) (4, 7, 14-21). We synthesized 4 of these SP peptides and 3 of the MGP peptides. Using the endogenous T cell (ETC) generation workflow (see Methods section), we generated individual T cell cultures against all 4 SP peptide and all 3 MGP peptides (Fig. 1). However, when these highly enriched (> 80% tetramer+) T cell cultures were tested against HLA-matched target cells engineered to express the relevant SARS-CoV-2 Spike protein or membrane glycoprotein, no evidence of target cell killing was observed (Fig. 2-8). We postulated that these *in silico* predicted peptides were not endogenously presented; and that a more accurate means of identifying endogenously presented, immunogenic epitopes would be desirable and could be achieved by directly eluting and sequencing peptides from the MHC of SARS-CoV-2-expressing cells.

**Fig. 1.**
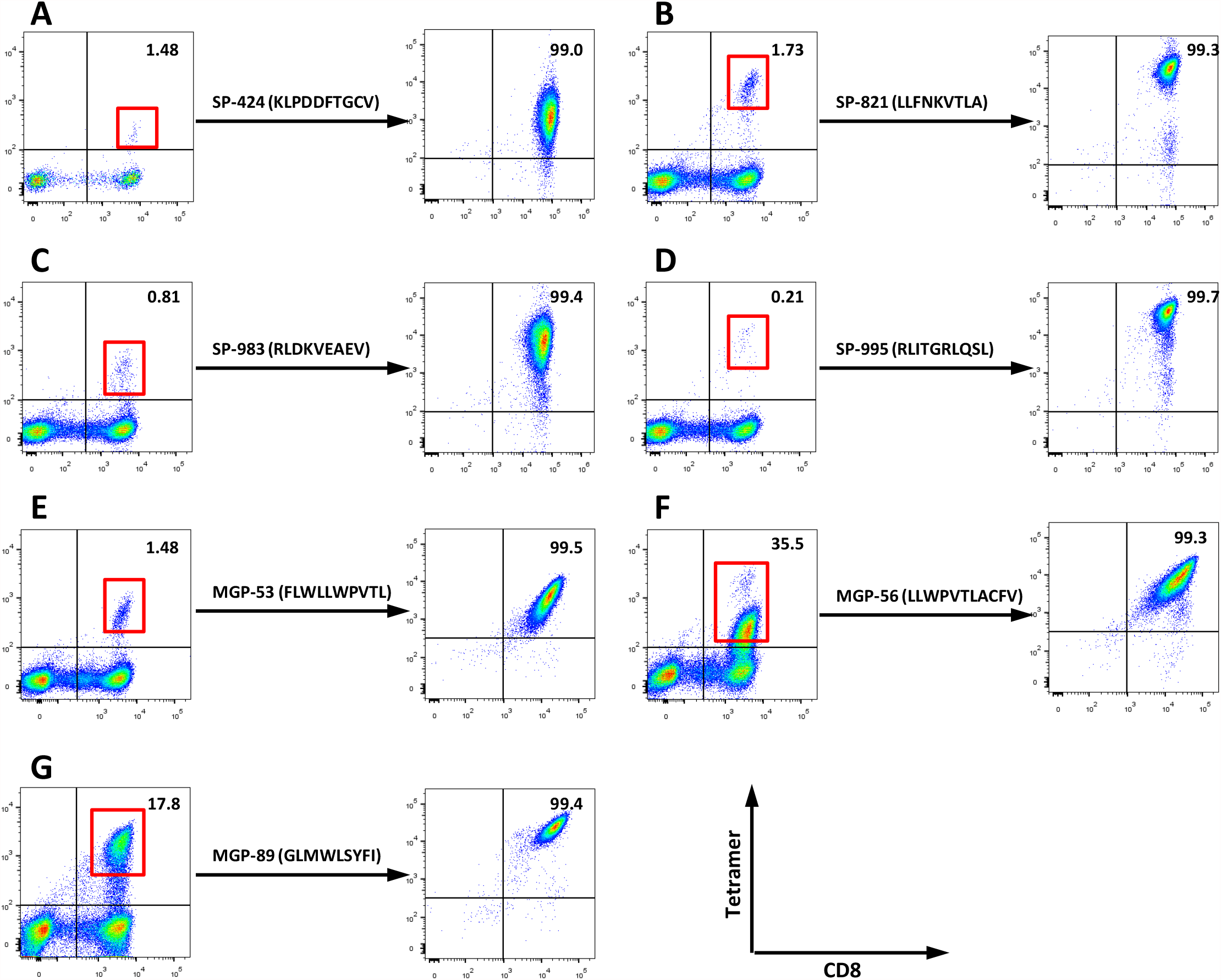
T cell generation for predicted SARS-CoV-2 HLA-A0201 restricted peptide. (A-D) 4 predicted spike protein (SP) HLA-A0201 peptide SP-424 (KLPDDFTGCV), SP-821 (LLFNKVTLA), SP-983 (RLDKVEAEV), SP-995 (RLITGRLQSL) and (E-G) 3 predicted membrane glycol-protein (MGP) HLA-A0201 peptides MGP-53 (FLWLLWPVTL), MGP-56 (LLWPVTLACFV), MGP-89 (GLMWLSYFI), were selected for antigen specific T cell generation using endogenous T cell (ETC) generation workflow. The peptide pulsed mature dendritic cells (DCs) were co-cultured with autologous PBMC from HLA-A0201+ healthy donors. After two rounds of stimulation, CD8+ and tetramer-positive T cells were induced (left side). After sorting and REP for CD8+ and Tetramer+ T cells with two weeks, high purity of specific CTLs were expanded (right side).

**Fig. 2.**
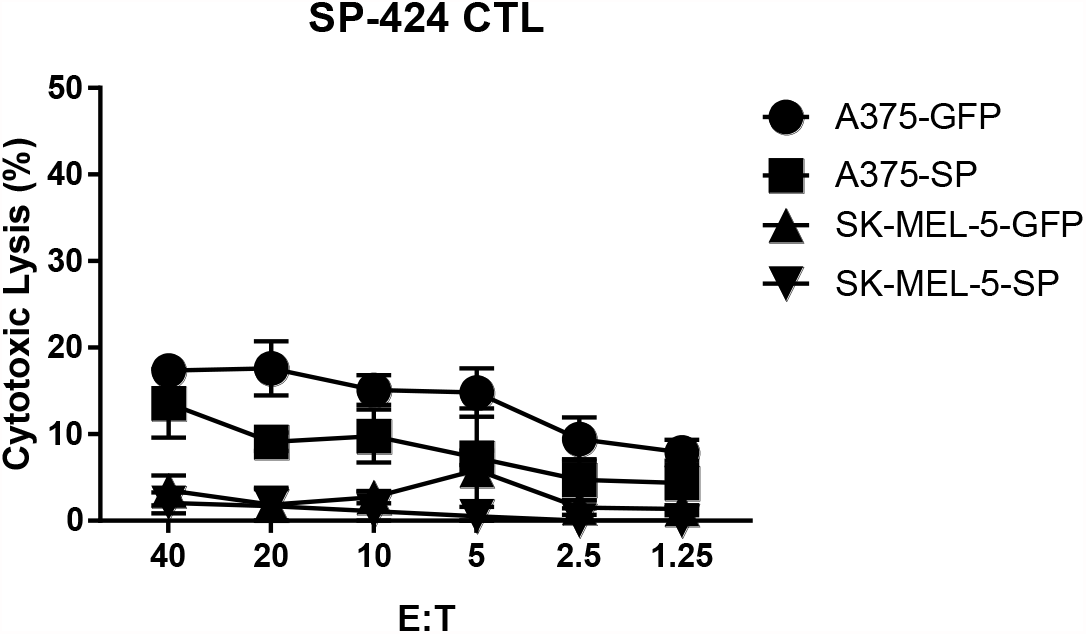
SP-424 CTL functional assay. The antigen specific-cytolysis were analyzed with standard ^51^Cr release assay (CRA) using SP or GFP force expressing HLA-A0201 cell lines (A375-SP, A375-GFP, SK-MEL-5-SP, SK-MEL-5-GFP) as targets.

**Fig. 3.**
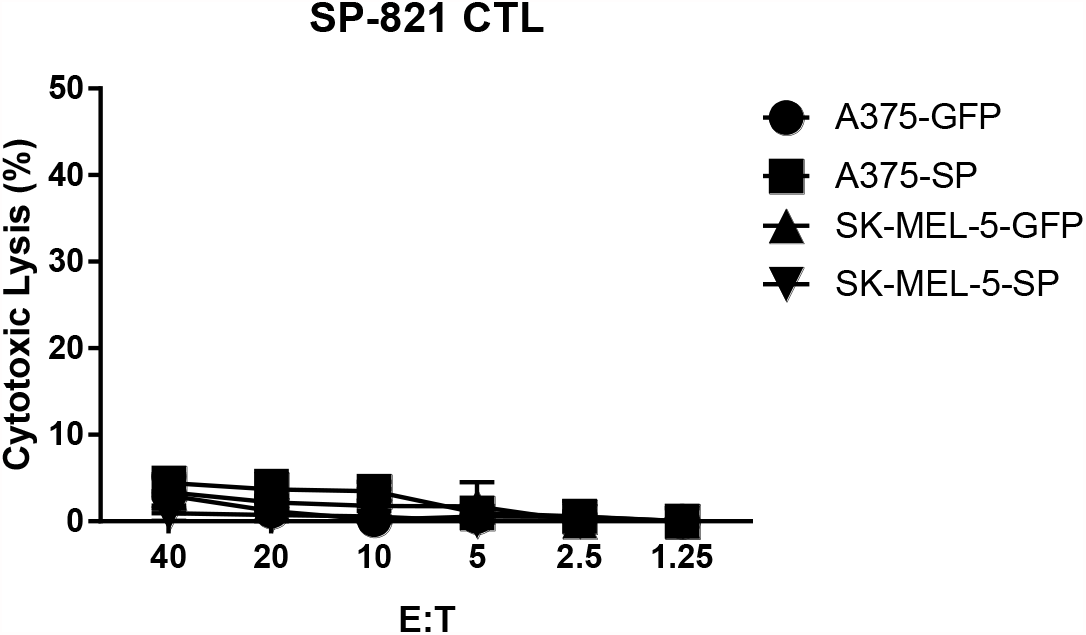
SP-821 CTL functional assay. The antigen specific-cytolysis were analyzed with standard ^51^Cr release assay (CRA) using SP or GFP force expressing HLA-A0201 cell lines (A375-SP, A375-GFP, SK-MEL-5-SP, SK-MEL-5-GFP) as targets.

**Fig. 4.**
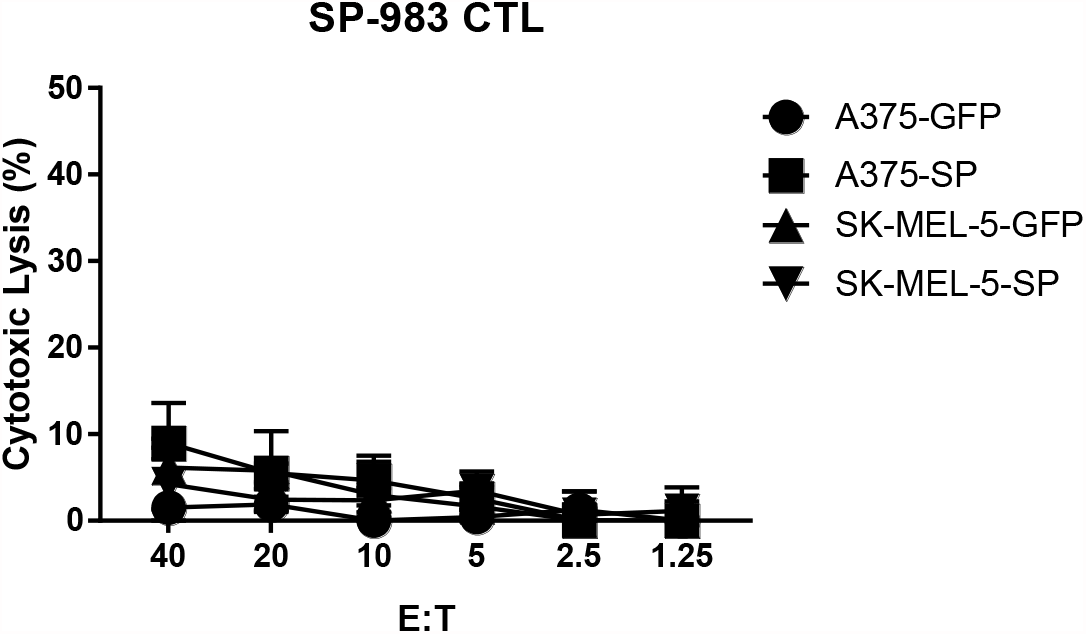
SP-983 CTL functional assay. The antigen specific-cytolysis were analyzed with standard ^51^Cr release assay (CRA) using SP or GFP force expressing HLA-A0201 cell lines (A375-SP, A375-GFP, SK-MEL-5-SP, SK-MEL-5-GFP) as targets.

**Fig. 5.**
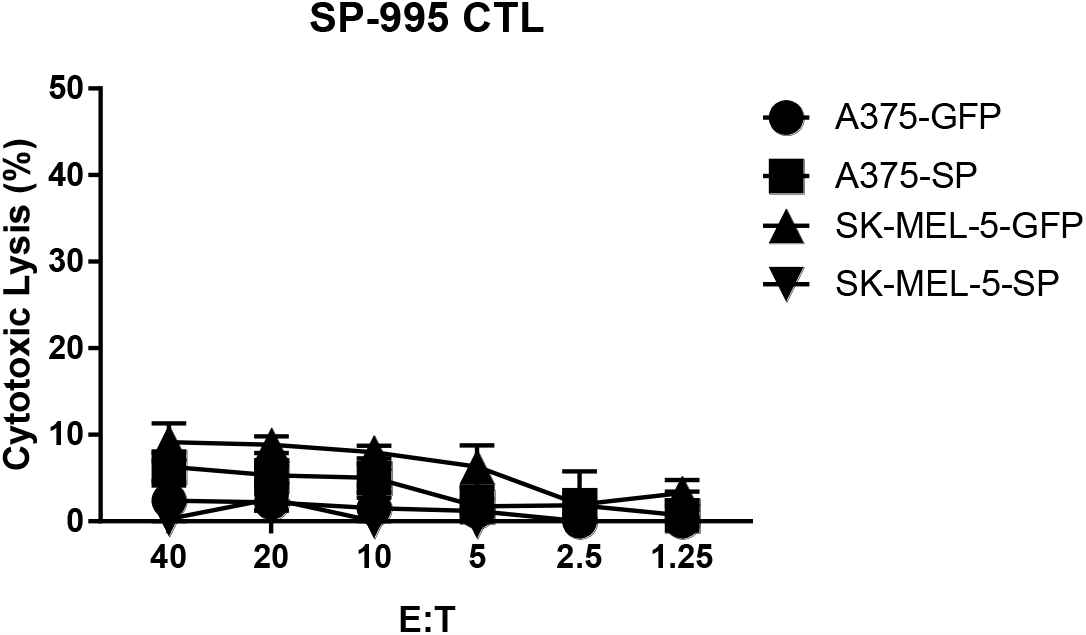
SP-995 CTL functional assay. The antigen specific-cytolysis were analyzed with standard ^51^Cr release assay (CRA) using SP or GFP force expressing HLA-A0201 cell lines (A375-SP, A375-GFP, SK-MEL-5-SP, SK-MEL-5-GFP) as targets.

**Fig. 6.**
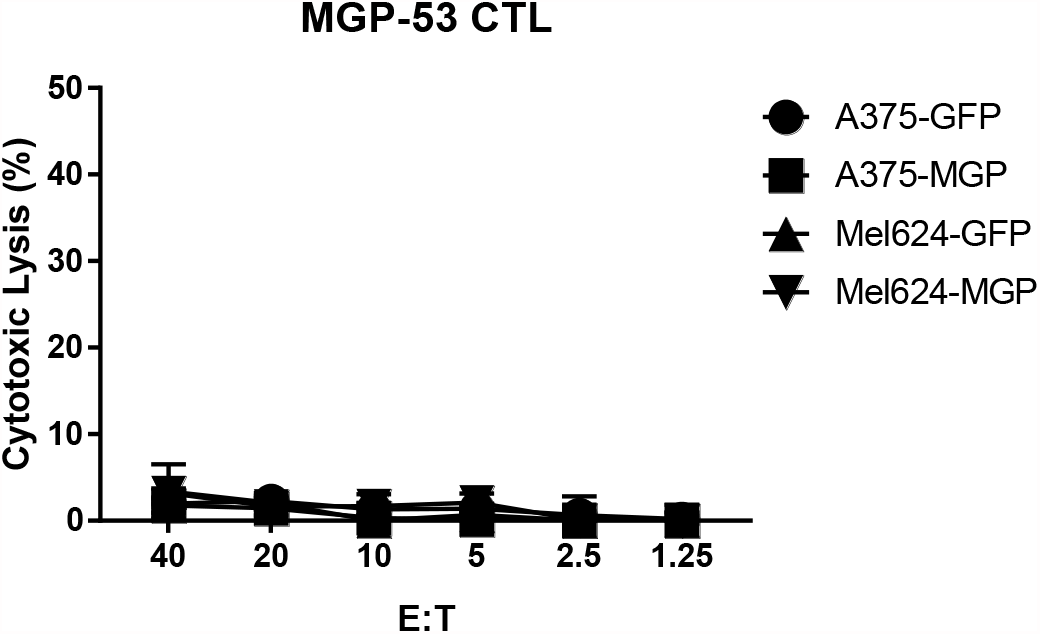
MGP-53 CTL functional assay. The antigen specific-cytolysis were analyzed with standard ^51^Cr release assay (CRA) using MGP or GFP force expressing HLA-A0201 cell lines (A375-MGP, A375-GFP, Mel624-MGP, MEL624-GFP) as targets.

**Fig. 7.**
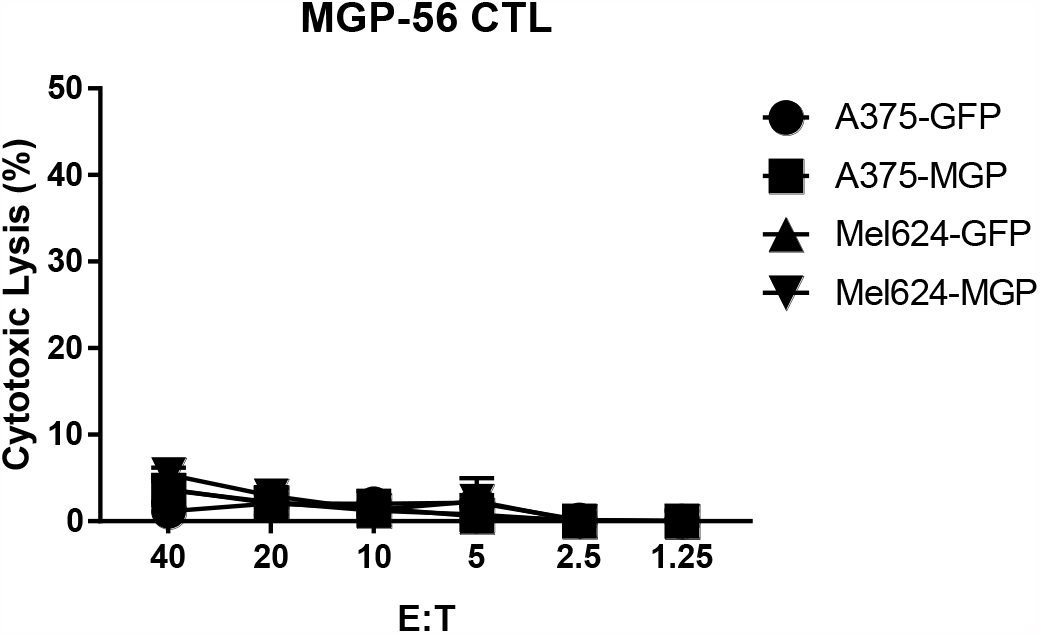
MGP-56 CTL functional assay. The antigen specific-cytolysis were analyzed with standard ^51^Cr release assay (CRA) using MGP or GFP force expressing HLA-A0201 cell lines (A375-MGP, A375-GFP, MEL624-MGP, MEL624-GFP) as targets.

**Fig. 8.**
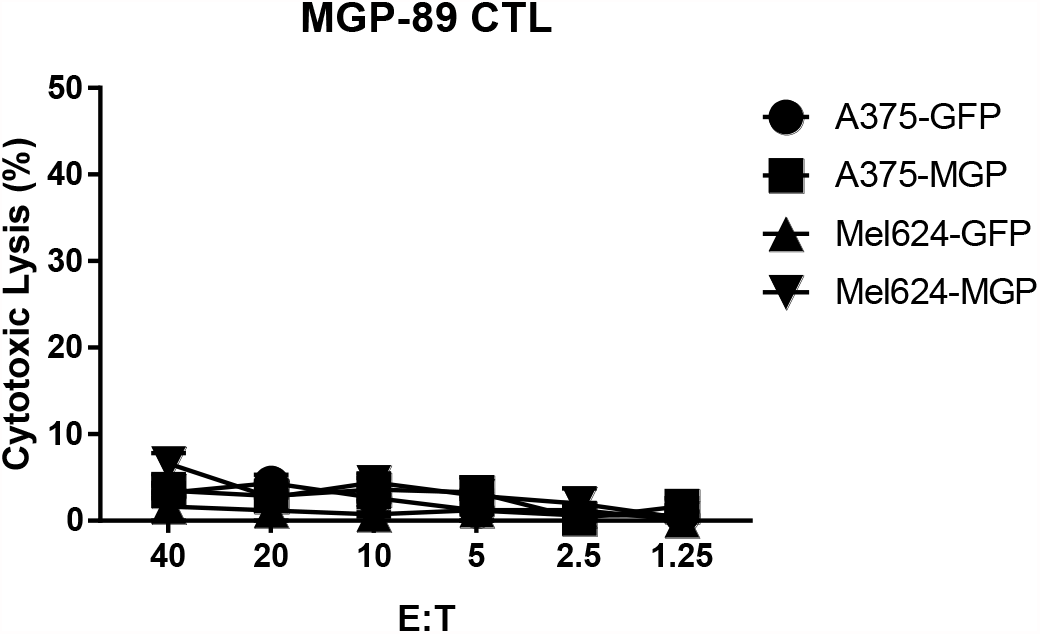
MGP-89 CTL functional assay. The antigen specific-cytolysis were analyzed with standard ^51^Cr release assay (CRA) using MGP or GFP force expressing HLA-A0201 cell lines (A375-MGP, A375-GFP, Mel624-MGP, MEL624-GFP) as targets.

## Discussion

To date, nearly 1,500 predicted Class I epitopes for SARS-CoV2 have been identified by in silico prediction methods, and in some cases, ‘validated’ by eliciting T cell responses using PBMC of patients with COVID19 (4, 5). These peptides have been used extensively to evaluate the T cell response of patients, and occasionally healthy donors, to COVID19, and COVID19 vaccines, and increasingly, to develop T cell-based therapies. What has not been demonstrated however, is whether any of these 1,500 predicted peptides are in fact processed and presented by SARS-CoV2+ cells and represent naturally-occurring epitopes recognized by T cells. A preliminary screen of predicted SARS-CoV2 epitopes considered “immunodominant” among widely cited reports appears to support this premise: we found that 7 of these 8 predicted peptides were unable to elicit a T cell response that would lead to recognition of SARS-CoV2+ targets suggesting that responses to these peptides may be artifactual, or at best cross-reactive (Fig. 1) (4, 7, 14-21). To date, there has been no empiric validation of SARS-CoV2 epitopes for immunogenicity.

Equally important in defining the landscape of COVID19 infection and control, its natural history, vaccine efficacy and therapeutic intervention, is an accurate measure of the SARS-CoV2-specific immune response. While the pools of predicted peptides currently in use to evaluate Class II and Class I -restricted responses have been used extensively and appear to provide a measure of overall immune response, SARS-CoV2-specific immunity is poorly defined when the majority of peptides may not be immunogenic. In contrast to in silico prediction methods, a strategy to more precisely and rigorously define immunogenicity would be desirable. Merely demonstrating that in silico predicted epitopes can elicit a peptide-specific T cell response from donor PBMC speaks only to the ability of a predicted peptide to bind MHC with high affinity and/or its ability to elicit cross-reactive T cells and fails to validate whether such in silico predicted peptides are presented or can elicit a bona fide SARs-CoV2-specific T cell response. Actual immunogenicity must be demonstrated by empiric fashion: i.e. such a peptide directly defined (e.g. by mass spectrometry of MHC peptidome) should be demonstrated to recognize target cells that endogenously express SARS-CoV2. A panel of such MS-defined peptides would be predicted to be more relevant when evaluating T cell immunity to COVID19 and when designing SARS-CoV2 vaccines.

## Materials and Methods

### Blood donors and Cell lines

Healthy donor peripheral blood mononuclear cells (PBMC) samples expressing the HLA-A0201 allele were purchased from HemaCare (CA, USA) as a source of responding T cells and autologous antigen presenting cells. TAP-deficient T-B cell hybrid cell line T2, melanoma cell line A375 (HLA-A0201), package cell line 293T were purchased from ATCC (VA, USA). Melanoma cell line SK-MEL-5 (HLA-A0201) were purchased from NCI. Melanoma cell line Mel624 (HLA-A0201) was the gift from Dr. Steven Rosenberg (NCI). Lymphoblastoid cell lines (LCL) are EBV-transformed lymphoblastoid cell lines established in our laboratory. Cancer cell lines were maintained in RPMI-1640 media with Hepes (25 mM), L-glutamine (4mM), penicillin (50 U/ml), streptomycin (50 mg/ml), sodium pyruvate (10 mM), nonessential amino acids (1 mM), and 10% fetal bovine serum (FBS) (Sigma, MO, USA).

### Lentivirus transduction

The cDNA of membrane glyco-protein (MGP) and Non-structure protein 13 (NSP13) of ORF1b from SARS-CoV-2 were purchased from Genscript (NJ, USA) and cloned into lentiviral vector pLVX (TAKARA, CA, USA) with fusion of GFP. In this vector, the expressing gene was driven by human EF1 promoter. MGP-pLVX or SP-pLVX lentiviral vector were transfected into package cell line 293T, together with package vectors contain VSVG envelop vector to make lentivirus. A375, SK-MEL-5, and Mel624 cell lines were infected with MGP-pLVX or SP-pLVX lentiviral vectors and the stable cell lines were screened with puromycin selection. MGP or SP gene expressing efficiency was detected by analyzing the percentage of GFP using flow cytometry (NovoCyte Flow Cytometer Systems, Agilent, CA, USA).

### Generation of SARS-CoV-2 specific T cells

Generation of antigen-specific T cell stimulation was performed according to our endogenous T cell (ETC) generation workflow (22). Briefly, adherent PBMCs were treated with GM-CSF (800 U/mL) and IL-4 (500 U/mL) for 6 days to generate immature DC (iDC), and the iDC were then matured with a cytokine cocktail containing TNF-α (10 ng/mL), IL-1β (2 ng/mL), IL-6 (1000 U/mL), PGE-2 (1000 ng/mL) for an additional 2 days. After 7 days in culture, T cell cultures were restimulated with peptide-pulsed DC as before. IL-2 (10 U/mL) and IL-7 (5 ng/mL) were added on the second day.

### Sorting and expansion

After two stimulation cycles, an aliquot of each well was stained with custom PE-conjugated MHC tetramer folded with HLA matched SARS-CoV2 peptide, and with APC-Cy7 conjugated anti-CD8 antibody (Biolegend, CA, USA). Cells were washed and analyzed by flow cytometry (NovoCyte Flow Cytometer Systems, Agilent, CA, USA). The tetramer positive staining wells were pooled and CD8/Tetramer double positive population were sorted using flow cytometric sorting (ARIA II sorter, BD, CA, USA) and then expanded using a rapid expansion protocol (REP) in a sterile 25 mL flask containing RPMI-1640 with Hepes (25 mM), L-glutamine (4mM), penicillin (50 U/ml), streptomycin (50 mg/ml), sodium pyruvate (10 mM), 10% fetal bovine serum (FBS), irradiated PBMC and LCL feeder cells, as previously described (23).

### Function analysis of SARS-CoV-2 specific T cells

The cytotoxicity of purified SARS-CoV-2 specific T cells following expansion was confirmed using standard chromium (^51^ Cr) release assay (CRA). Peptide dose titration experiments were performed to test cognate peptide recognition of SARS-CoV-2 CTL.

## Statistical analysis

Data analysis was performed using GraphPad Prism version 7.03. Normally distributed data were analyzed using parametric tests (ANOVA or unpaired t test). All *p* values are indicated in figure legends. Flow Cytometry data were analyzed using FlowJo (version 10).

## Acknowledgments

We would like to thank the help from South Campus Flow Cytometry & Cell Sorting Core of UT MD Anderson Cancer Center, which is supported by NCI P30CA016672. This work was supported by National Institutes of Health grant RO1 3R01CA237672-02S1 (to C.Y.) and the Parker Institute of Cancer Immunotherapy (C.Y.)

## Author Contributions

K.P., Y.C., C.Y. conducted overall experimental design.K.P. established SARS-CoV-2 MGP and SP overexpressing cell lines. K.P., M.C., J.W., I.L., R.S., S.S. conducted SARS-CoV-2 MGP and SP specific CTL generation and functional validation. K.P., Y.C., S.S., C.Y. wrote the paper. C.Y. was responsible for the supervision of laboratory studies and manuscript writing and editing.

## Competing Interest Statement

C.Y. serves as a member for Parker Institute for Cancer Immunotherapy. All other authors declare that they have no competing interests.

